# Improving isoform-level eQTL and integrative genetic analyses of breast cancer risk with long-read RNA transcript assemblies

**DOI:** 10.64898/2026.03.22.713514

**Authors:** S. Taylor Head, Aryun Nemani, Yung-Han Chang, Tabitha A. Harrison, Sean T. Bresnahan, Joseph H. Rothstein, Weiva Sieh, Sara Lindström, Arjun Bhattacharya

## Abstract

Most eQTL and TWAS analyses quantify expression using aggregate, tissue-agnostic transcript annotations and ignore isoform-level regulation, potentially obscuring or misattributing regulatory mechanisms. Here, we developed a framework leveraging publicly available long-read RNA-seq data to perform tissue-informed inference of genetic regulation and prioritize candidate causal isoforms for breast cancer risk. We quantified gene- and isoform-level expression in breast tumor (TCGA), non-cancerous mammary tissue, and cultured fibroblasts (GTEx) using three transcriptome annotations: standard GENCODE, tissue-specific long-read-derived assemblies, and combined annotations incorporating transcript-isoforms from both. While GENCODE cataloged over 250,000 pan-tissue isoforms, the tissue-specific long-read assemblies captured reduced sets of 74,717 isoforms in tumor, 48,057 in fibroblasts, and 22,941 in healthy breast. We performed eQTL mapping and fine-mapping, followed by colocalization with overall and subtype-specific breast cancer GWAS and isoform-level TWAS. While most eGenes were concordant across annotations, approximately 1/3 of lead cis-eQTLs for shared eGenes differed between long-read assemblies and GENCODE. Further, eIsoform discovery was highly annotation-specific. In healthy breast tissue, the gold standard tissue for building gene expression prediction models for TWAS of breast cancer, 46% of eIsoforms identified by the long-read annotation were unique to that annotation even though 93.7% of them are present in GENCODE. Despite combined annotations expanding the GENCODE catalog by only 0.6-7.6% depending on tissue source, 69% of unique significant isoform-trait associations were specific to a single annotation. Long-read-informed annotations uncovered regulatory associations entirely missed by GENCODE, including a candidate regulatory isoform at the *MARK1* locus captured only in fibroblasts and a previously unannotated splice variant prioritized as the likely effector transcript at *NUP107*. These findings demonstrate that transcript annotation is not merely a technical consideration but critically defines the biological hypothesis space for regulatory mechanisms and shapes discovery. Incorporating tissue-resolved isoform annotations from long-read RNA-seq improves the specificity of regulatory inference and enhances identification of candidate causal isoforms at GWAS loci.

## INTRODUCTION

Identifying the likely transcriptomic mechanisms affected by the >200 genetic variants linked to breast cancer risk remains an open challenge^1–7^. While integrative approaches such as colocalization and transcriptome-wide association studies (TWAS) help identify causal genes at breast cancer genome-wide association study (GWAS) loci, these approaches often rely on total gene expression aggregated across transcript isoforms, potentially obscuring isoform-specific regulatory effects. Indeed, modeling isoforms rather than genes has been found to increase the discovery of transcriptome-mediated genetic associations with complex traits by as much as 60%^8–10^. Ignoring isoforms is a notable limitation in breast cancer, where isoform-specific antagonistic effects are well-documented in genes such as *BRCA1*, *ERBB2*, *KLF6*, *AIB1*, *TNC37-42*, and *ESR1*^11–16^.

Both gene- and isoform-level integrative analyses depend upon accurate quantification of expression. Aggregate transcriptome references like GENCODE, which are often used for short-read RNA-seq quantification, can obscure the tissue-specific regulatory mechanisms underlying tumorigenesis. They can contain hundreds of thousands of annotated isoforms, many of which are only weakly or not at all expressed in breast tissue^17,18^. Isoform quantification is further complicated by the limitations of short-read RNA-seq approaches that can struggle to accurately resolve transcript isoforms because of read-to-transcript mapping ambiguity^19^. When sequenced fragments align to exons shared by multiple isoforms, which is a frequent occurrence in genes with complex splicing patterns, probabilistic assignment of reads generates substantial uncertainty in isoform-level expression estimates^20^, leading to reduced statistical power in downstream integrated analyses^19^. However, recent technological advances in long-read RNA-seq have enabled more comprehensive detection of full-length transcript isoforms. While short-read sequencing typically produces reads under 300 bp in length, long-read RNA sequencing technologies can generate reads up to 15-20 kilobases that span entire transcripts. This offers direct insight into exon structure and a more comprehensive view of transcript isoform diversity within a tissue or cell type^21–25^.

In this study, we leveraged long-read RNA-seq-derived transcriptome references for breast-relevant tissues to deliver an improved, tissue-specific view of genetic regulation at the isoform level across non-cancerous mammary tissue, malignant breast tumors, and cultured fibroblasts. We compared gene- and isoform-level expression quantitative trait loci (eQTL) mapping, colocalization, and TWAS for overall and subtype-specific breast cancer across multiple annotations and tissues. Our analyses yielded several major findings. First, although the long-read transcriptome assemblies are highly specific and contain >100-200K fewer transcripts than GENCODE, they retained comparable power for mapping GWAS loci with detected eQTLs, indicating robust coverage of risk-associated loci. Second, the long-read-informed analyses revealed multiple novel isoform-level regulatory mechanisms that are undetectable with standard reference transcriptomes. Third, we demonstrated that colocalization and TWAS results are highly sensitive to even minor changes in the transcriptome assembly used to quantify short-read RNA-seq samples, underscoring the importance of reference selection for downstream analyses. Finally, we provide an initial catalog of germline genetic effects on isoform-level expression across tissue states that can serve as a resource for future studies of breast cancer.

## RESULTS

In this study, we re-quantified short-read RNA-seq data from non-cancerous breast tissue, breast tumors, and fibroblasts using long-read derived transcriptome assemblies. We then systematically compared *cis*-eQTL mapping, colocalization of eQTL and breast cancer GWAS signals, and isoform-level TWAS results against those obtained using GENCODEv45 and a combined annotation (Fig. 1a). Despite containing 3-11 times fewer transcripts than GENCODE, long-read assemblies recovered comparable numbers of eQTL-linked independent GWAS loci. However, they yielded markedly fewer colocalized loci and TWAS associations, potentially reflecting tissue specificity of causal regulatory mechanisms and reduced spurious findings rather than diminished power. Aggregate annotations like GENCODE can dilute signal across structurally similar transcripts and misattribute reads to transcripts that are lowly or not expressed in the target tissue, elevating quantification uncertainty and producing unstable colocalization or TWAS results (Fig. 1b). Here, we show how long-read assemblies can help resolve isoform-level ambiguity at known GWAS loci, uncover novel isoforms mediating genetic risk, and inform biological interpretation of potential disease mechanisms in the target tissue context.

**Figure 1.**
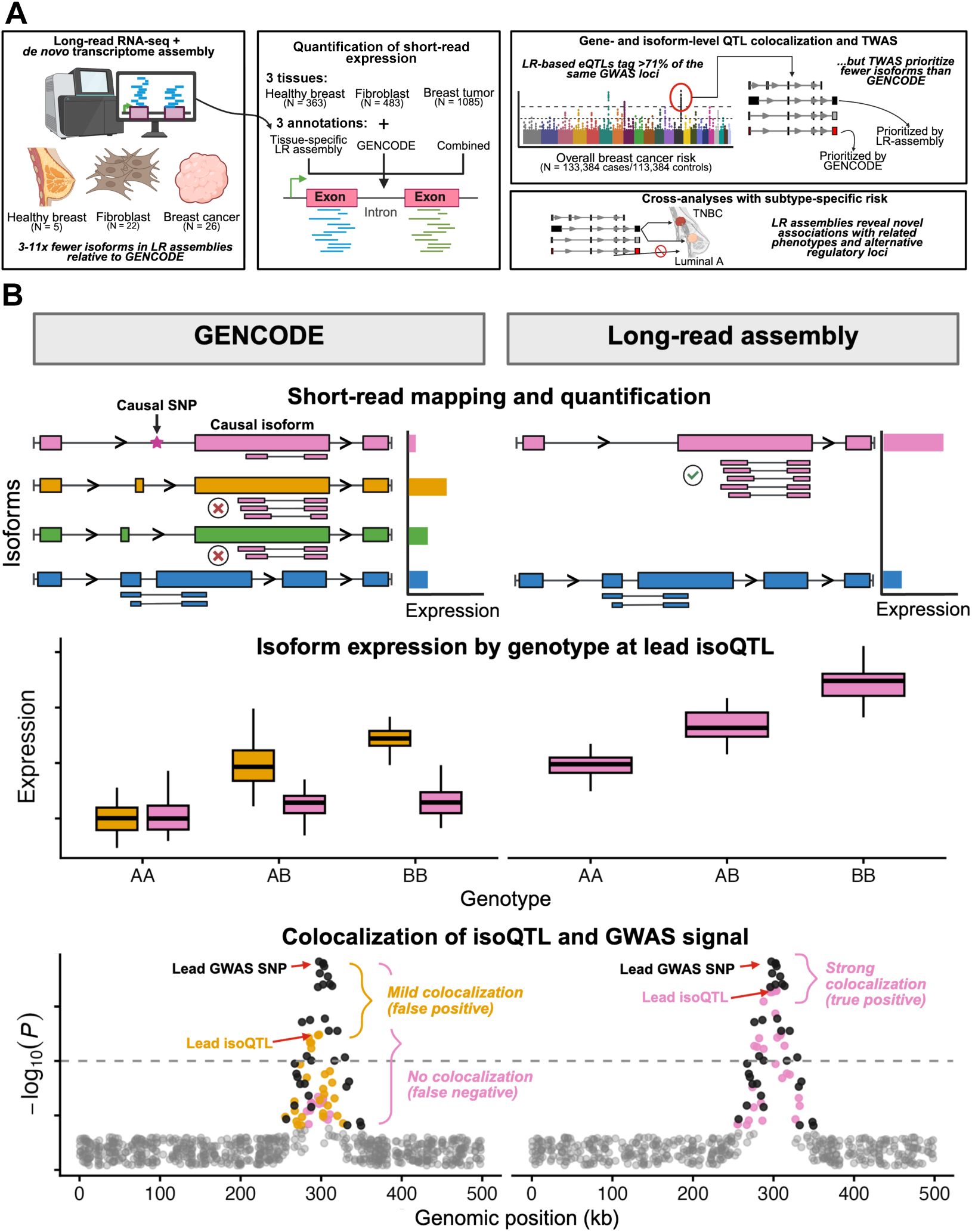
**(A)** In this project, we developed three de novo transcriptome assemblies in healthy breast tissue, fibroblasts, and breast tumor tissue derived from publicly available LR RNA-seq data (left panel). Next, we used these tissue-specific LR assemblies, the GENCODEv45 reference transcriptome, and combined annotations each to re-quantify short-read RNA-seq samples in healthy breast tissue, fibroblasts, and breast cancer tumor tissue (middle panel). We performed isoform- and gene-level eQTL mapping and fine-mapping, as well as colocalization and TWAS analyses with overall breast cancer GWAS risk loci and subtype-specific breast cancer loci (right panel). (**B**) In this example, the GENCODE transcript reference annotation contains four structurally similar isoforms for a gene. Only two are relevant to the tissue of interest and only one is causal for a trait, leading to misattribution of reads to irrelevant isoforms in GENCODE. This can propagate downstream and manifest as inaccurate isoform expression quantification, spurious eQTL mapping, and false positives and negatives in colocalization with GWAS loci. Abbreviations: LR, long-read; eQTL, expression quantitative trait loci; TWAS, transcriptome-wide association study; isoQTL, isoform-level eQTL; GWAS, genome-wide association study.

### Long-read transcriptome assemblies greatly refine gene and isoform space

We applied a robust transcriptome assembly and quantification framework^26,27^ to jointly discover and quantify expression of transcript isoforms in three sets of breast cancer-relevant long-read RNA-seq libraries (Fig. 1a). Compared to the GENCODE reference transcriptome containing 252,931 transcript isoforms across 63,187 genes^28^, these *de novo* long-read assemblies yielded markedly lower numbers of annotated isoforms and genes, both before and following comprehensive quality control protocols. Across the three tissues, isoform counts from long-read assemblies decreased over 70% relative to GENCODE, and this corresponded to a decrease in gene counts by over 63% (Fig. 2a). We resolved the greatest number of high-confidence isoforms (74,717) in our malignant breast tumor samples (N=26; primary biopsies, patient-derived xenograft tumors, and cell lines) mapping to 18,100 genes. The healthy breast samples (N=5; primary mammary tissue and cell lines) demonstrated the most pronounced transcriptome refinement, with only 22,941 high-confidence isoforms assembled across 11,015 genes.

**Figure 2.**
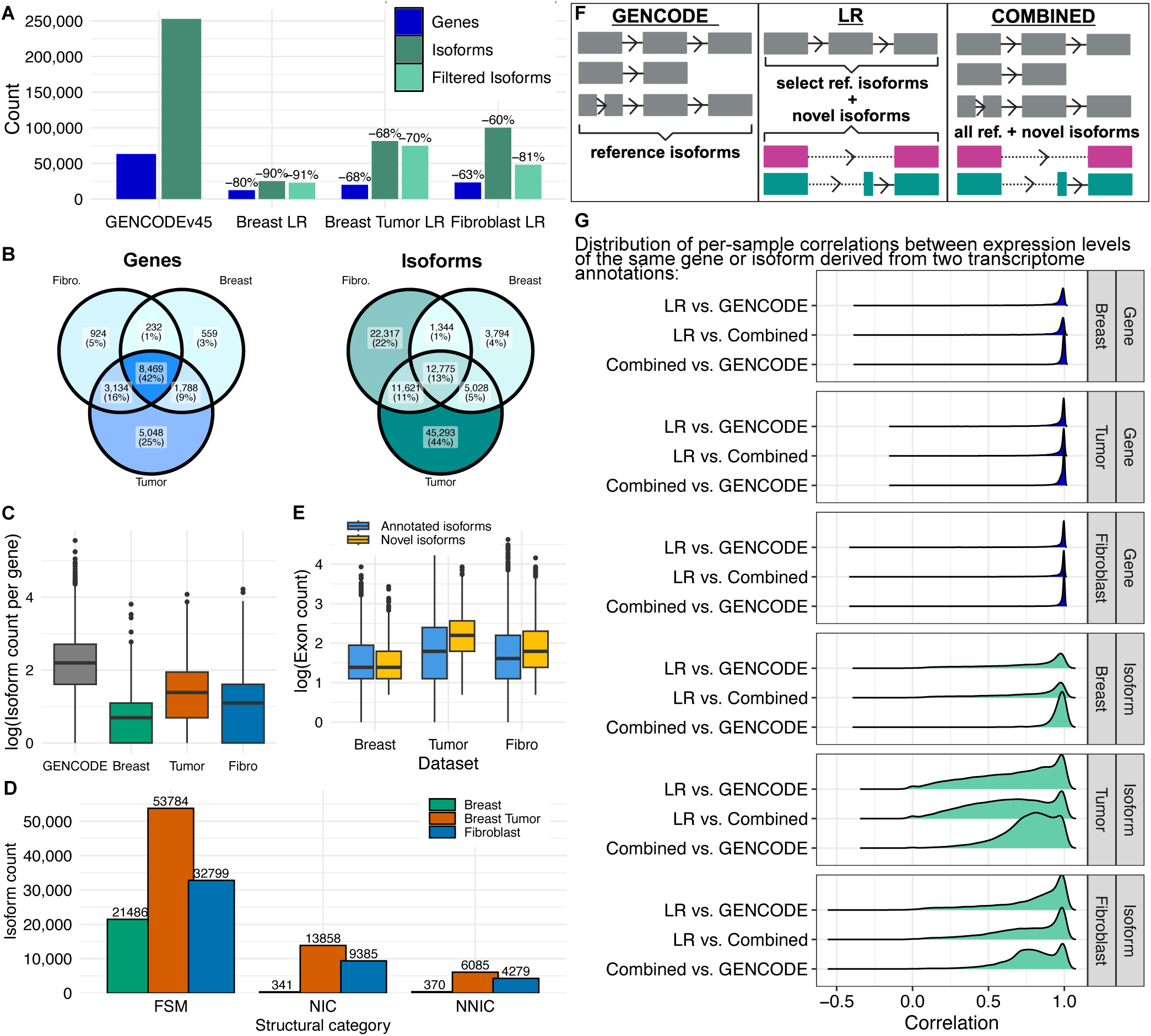
(**A**) Number of genes and isoform transcripts in the GENCODEv45 reference transcriptome compared to LR RNA-seq-derived healthy breast, breast tumor, and fibroblast transcriptome assemblies. Percentages shown represent percent decrease in these counts relative to GENCODE. (**B**) Overlap in post-filtering genes and isoforms across the three *de novo* long-read assemblies. (**C**) Log isoform counts per gene in LR derived assemblies and GENCODE for genes present in both long-read assemblies and GENCODE reference. Whiskers of boxplots extend to maximum/minimum point that is less than 1.5 × interquartile range from the third/first quartiles. **(D**) For each LR assembly, distribution of log exon counts for novel, unannotated isoforms versus transcripts present in GENCODE. (E) High confidence transcript isoforms assembled in the LR RNA-seq derived stratified by transcript structural category. (**F**) Graphical representation of how the three transcriptome annotations were constructed in each of our three short-read tissue contexts. Combined annotations incorporate all isoforms present in GENCODEv45 appended with all novel, non-FSM isoforms assembled in the long-read read samples of a given tissue. (**G**) Correlation of expression levels in re-quantified short-read samples across annotations for each gene (blue) and isoform (green) present in each pairwise annotation set. Abbreviations: LR, long-read; FSM, full splice match isoforms to annotated transcript; NIC, novel in catalog isoform; NNC, novel not in catalog isoform.

Given the limited number of healthy breast libraries, cell-cultured fibroblasts provided an additional non-malignant and non-epithelial reference tissue with a long-read sample size (N=22) comparable to breast tumor. We observed filtering of approximately 91% of isoforms in both healthy breast and tumor. However, in fibroblasts, we observed the most stringent quality control filtering of long-read isoform calls, with 48% (48,057 isoforms of 100,190 assembled) mapped to a total of 12,990 genes retained as high-confidence. 92.3% of all filtered isoforms lacked sufficient splice junction coverage in short-reads or full-length reads (Fig. S1, Table S1).

While we consistently detected 8,469 shared genes and 12,775 shared isoforms across all three long-read assemblies (Fig. 2b), 58% of genes and 87% of long-read isoforms displayed some degree of tissue-specificity. Differences in detected and retained tissue-specific transcriptome sizes likely reflect a combination of tissue-specific alternative splicing programs, long-read sample size, and heterogeneity of long-read sample type. In particular, there are 5,048 genes and 45,293 isoforms unique to the breast tumor assembly, comprising 25% and 44% of all assembled features (Fig. 2b). For ubiquitous genes, the number of isoforms per gene was substantially lower than in GENCODE (median 9) across all assemblies, but the tumor assembly again retained the highest number (median 4; Fig. 2c). Overall, isoform counts per gene decreased relative to the reference for 78.3%, 81.4%, and 91.1% of shared genes in tumor, fibroblast, and healthy breast long-read assemblies, respectively (Fig. S2). These observations of increased isoform diversity in the tumor assembly are consistent with abnormal regulation of alternative splicing in malignant tumors relative to healthy tissue^29–31^.

### Long-read assemblies reveal novel isoform structures in breast tumor and fibroblasts

The majority of long-read-derived isoforms are full-splice matches (FSM) with splice junction structure perfectly matching an annotated isoform in the reference GENCODE transcriptome: 68.3%, 72%, and 93.7% in fibroblasts, tumor, and healthy breast, respectively (Fig. 2d). However, there was a non-negligible contribution of novel transcript isoform structures to the tumor and fibroblast assemblies (Table S2). While there were only 1,455 novel isoforms in healthy breast tissue, we assembled 20,933 novel isoform structures in tumor tissue and 15,258 in fibroblasts. The vast majority (66.2% and 61.5%) of these were “novel in catalog” (NIC) isoforms, reflecting a new combination of previously annotated splice junctions, and 28-29% were “novel not in catalog” (NNIC) isoforms containing one or more previously unannotated splice donor or acceptor sites^27^. Novel tumor isoforms also contained more exons (median 9 [IQR 7]) than their corresponding FSM counterparts (median 6 [IQR 8]; Fig. 2e) and were significantly longer (one-sided t-test P<2.2E-16). While these findings highlight the potential of long-read transcriptome assemblies to capture complex tissue-specific isoform structures currently missing from aggregate references, we note that the higher number of novel isoforms in tumor and fibroblasts may reflect a combination of biological differences and technical factors including varied sample sizes and sample composition (tissue vs. cultured cells).

Importantly, using these assemblies as refined transcriptome annotations directly impacted expression quantification in our short-read RNA-seq datasets (N=363 healthy breast mammary tissue, N=483 cell-cultured fibroblasts, N=1,085 breast tumors). We used GENCODE and the *de novo* long-read assemblies to construct three annotation sets per tissue (Fig. 2f): GENCODE isoforms alone, tissue-matched long-read assemblies alone (“LR”), and all GENCODE isoforms plus novel high-confidence non-FSM isoforms identified in the corresponding long-read samples (“combined”). Per-sample correlations of expression estimates across annotation pairs revealed that gene-level quantification is relatively robust to annotation choice, with consistently high correlations across all tissues (Fig. 2g). Isoform-level correlations, however, were markedly more variable. They exhibited heavy-tailed distributions even between minimally divergent catalogs like GENCODE and the combined annotations, despite the latter containing only 0.6-7.6% more isoforms. In practical terms, a low correlation here means that the relative expression of an isoform across samples is not preserved between annotations and suggests isoform-level quantification is highly sensitive to the transcriptome annotation choice, consistent with prior studies^32,33^.

### Long-read transcriptomes reveal alternative genetic architectures for shared regulatory features

To identify regulatory genetic variants that may underlie causal breast cancer pathways, we first performed eQTL mapping in each tissue. We identified 5,827, 2,305, and 1,177 genes in fibroblasts, breast, and breast tumor, respectively, as harboring one or more significant eQTL (eGenes) that replicated across all three annotations (Fig. 3a, Online Data). These common eGenes represented over half (52.9%) of all fibroblast eGenes (identified by any annotation), 31.3% of all breast eGenes, but less than 18% of tumor eGenes identified by at least one annotation.

**Figure 3.**
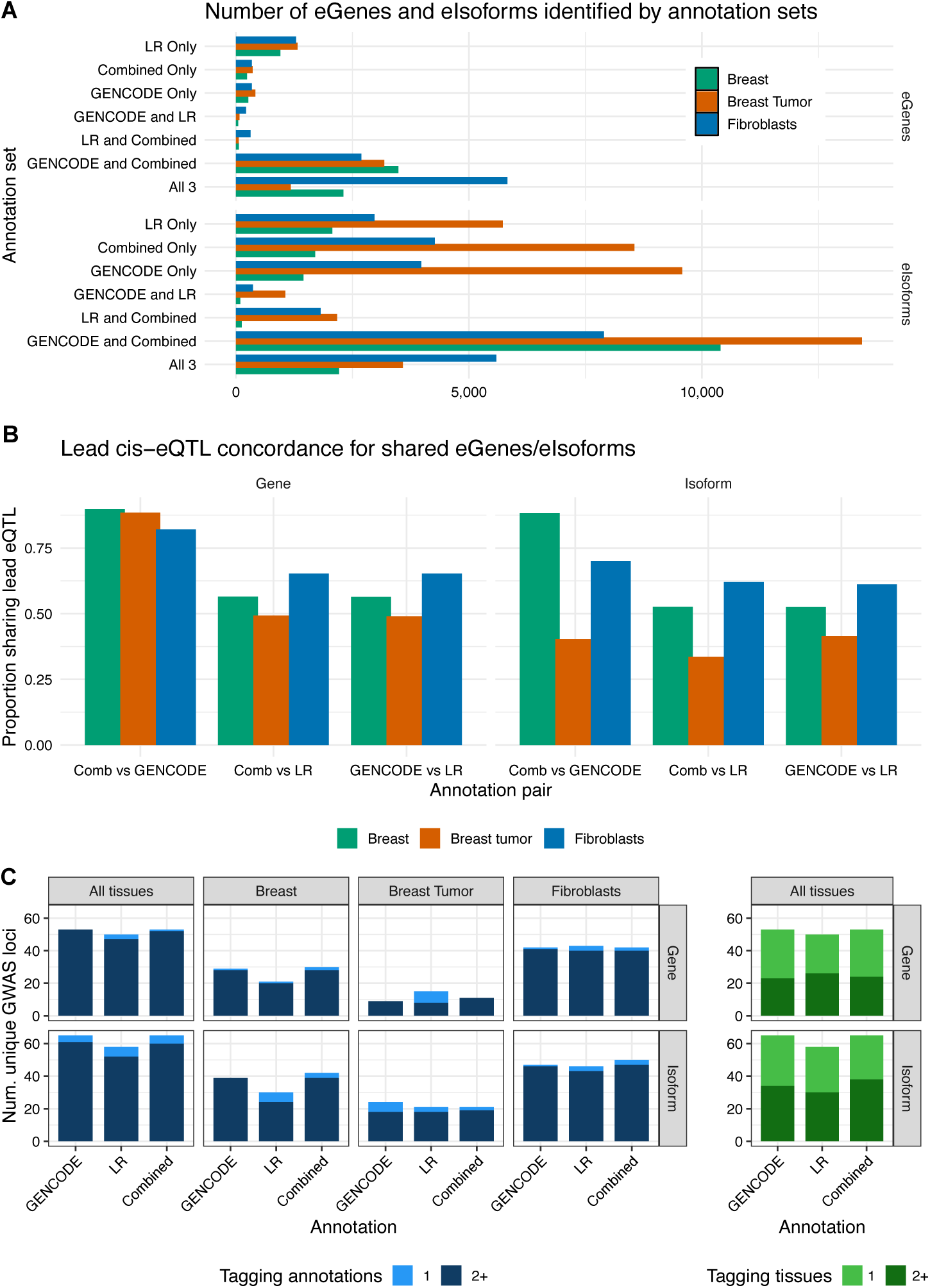
(**A**) Bars indicate the overlap of eIsoforms (isoforms with at least one significant eQTL) and eGenes (genes with at least one significant eQTL) identified in short-read healthy breast, breast cancer tumors, and fibroblasts across the three annotations. Sets representing intersections of two annotations (e.g., GENCODE and LR) indicate eGenes/eIsoforms identified only in those two annotations and not the third. (**B**) Bar heights reflect the proportion of eGenes (left) and eIsoforms (right) sharing the same lead cis-eQTL for features shared across annotations. **(C**) Gene- and isoform-level eQTL tagging of independent breast cancer GWAS loci. A tagged locus indicates an eQTL is within 1Mb and LD R^2^>0.9 with the GWAS risk variant. The bars are colored to reflect the proportion of GWAS loci that were uniquely tagged by that single annotation (left) or in a single tissue (right). For all panels (**A**)-(**C**), eQTLs were identified from the conditional linear model analysis scheme. Abbreviations: LR, long-read; eQTL, expression quantitative trait loci; LD, linkage disequilibrium; GWAS, genome-wide association study; comb, combined.

As expected, GENCODE and combined annotations showed particularly strong concordance in eGenes, with over 89% of eGenes overlapping between the two in each tissue. More notably, long-read assemblies consistently identified a substantial number of unique features not detected by either reference-inclusive annotation: 1,289 unique eGenes in fibroblasts, 954 in healthy breast, and 1,320 in tumor, accounting for 16.9%, 28.3% and 50.2% of all eGenes identified by long-read in each tissue. Fine-mapped eQTLs yielded similar patterns of cross-tissue and -annotation eGene discovery (Fig. S3).

While the proportion of long-read unique eIsoforms identified in tumors and in fibroblasts did not differ considerably from that observed for eGenes (differing only by 4.5-10.8%, Fig. 3a), healthy breast tissue showed a striking departure. Approximately 46.0% of all eIsoforms identified in healthy breast tissue using the long-read annotation were unique hits not similarly identified with GENCODE or combined annotations. This represents a 1.6-fold increase relative to the gene-level rate of long-read-specific signals. Notably, this occurred despite 93.7% of all isoforms in the healthy breast long-read assembly being FSM to previously annotated transcripts.

We also observed substantial annotation-specific eIsoform signals for reference-informed annotations. For example, in breast tumor, 34.6% of GENCODE eIsoforms were unique despite all being present in the combined tumor transcriptome annotation file, and this jumped to 43.7% following fine-mapping. These findings further suggest that isoform-level regulatory architecture is far more sensitive to transcriptome annotation choice than gene-level analyses, and while the degree of sensitivity differs across tissues, long-read-specific signals persist across all three tissue contexts. We provide full lists of eGenes and eIsoforms from conditional linear modeling and fine-mapping online (see Online Data).

We next compared the underlying regulatory variants driving shared eGenes and eIsoforms across annotation sets. Lead cis-eQTLs for shared genes were highly concordant between the GENCODE and combined annotations (>82.1%) but substantially less-so for comparisons involving the long-read assemblies (49.0-65.3%, Fig. 3b). This pattern persisted at the isoform level in fibroblast and healthy breast tissue, yet concordance dropped markedly in tumor where fewer than 41.5% of shared eIsoforms had matching lead isoQTLs across any annotation pair, including GENCODE-informed.

Among the shared regulatory features (eGenes and eIsoforms) with discordant lead eQTLs, linkage disequilibrium (LD) R^2^ was often moderate to high (median R^2^=0.83) but highly variable (IQR=0.59). This suggests that eQTL discordance was not solely driven by the selection of near-identical proxy variants (Fig. S4). Expanding beyond the single top variant to all significant cis-eQTLs per feature, we saw similar patterns of discordance across annotation sets. (Fig. S5).

While the eGenes and eIsoforms identified above through linear regression modeling have strong genetic signals, standard eQTL mapping does not confidently resolve the underlying causal variants. To address this, we performed Bayesian eQTL fine-mapping to more rigorously compare the sets of regulatory features identified across annotations while more directly incorporating LD structure. These fine-mapping results further reinforced observed discordance. Although the number of credible sets per feature and their sizes were comparable across annotations, the composition of these 95% coverage credible sets differed substantially. Among shared eGenes and eIsoforms identified by both GENCODE and long-read annotations, 20.0-31.2% showed <50% overlap in credible set variants (Fig. S6). Even at the stage of eQTL mapping, annotation choice can lead to materially different inferences about putative causal regulatory variants.

### eQTLs in long-read assemblies tag GWAS loci at similar rates to GENCODE

Strikingly, despite the reduced gene and isoform space cataloged by each long-read assembly, long-read-derived eQTLs tagged 332 independent breast cancer GWAS risk loci in tumor and fibroblasts at rates comparable to those observed for the larger GENCODE-inclusive annotations (Fig. 3c). Across annotations, between 41.5-56.6% of all tagged loci were observed in one tissue only, suggesting substantial tissue-specific regulatory architecture.

When aggregating across all tissues, long-read-defined eQTLs tagged >89.2% as many loci as GENCODE or combined annotations. Only in healthy breast tissue did the long-read annotation tag fewer loci overall compared to reference-inclusive annotations. However, the number of loci tagged by long-read eQTLs still exceeded 72.4% of the number tagged using GENCODE for both eGenes and eIsoforms. This is notable, given the breast long-read assembly contains 80% fewer genes and 91% fewer transcripts than GENCODE. These patterns remain largely consistent across different LD thresholds used to define locus tagging and restriction to fine-mapped eQTLs (Fig. S7).

With the exception of gene-level eQTLs in breast tumor, where 46.7% of loci tagged by long-read were unique, fewer than 20.0% of loci tagged by long-read were unique across remaining tissues. This indicates that long-read annotations largely implicated the same GWAS loci as reference transcriptomes. However, the specific eGenes and eIsoforms underlying these shared loci differed substantially. Fibroblasts provide an example, as this tissue showed the greatest number of tagged GWAS loci and similar tagging rates across all annotations. Among shared GWAS loci in fibroblasts, eGenes were nearly perfectly concordant between GENCODE and combined annotations, whereas concordance dropped to an average of 72.9% at the isoform level (Fig. S8). Concordance was lower still when comparing long-read to GENCODE, with only 42.1% average overlap in eIsoforms tagging the same shared locus.

Together, these results suggest that expansive reference annotations do not necessarily yield expanded insight into trait-relevant regulation. Long-read assembly-derived eQTLs tagged GWAS loci at rates largely similar to those observed with GENCODE. While most of these loci were consistent across annotations, this did not imply concordance in the underlying regulatory features. Instead, the specific eGenes and eIsoforms implicated at shared loci differed depending on the transcriptome annotation used for expression quantification. Signals relevant to breast cancer risk thus appear to be captured well by a subset of tissue-relevant isoforms reliably observed with long-read RNA-seq. These findings suggest that long-read assemblies provide a more parsimonious and tissue-informed representation of the regulatory landscape while retaining the capacity to implicate genetic loci relevant to breast cancer risk.

### Transcriptome annotation drives colocalization and TWAS hits

To further evaluate how transcript annotation shapes inference at GWAS loci, we compared isoform-trait associations prioritized by both colocalization and TWAS for risk of breast cancer (overall and five intrinsic-like subtypes). Across annotations, we identified 556 unique significant associations, each defined by an isoform, trait, and tissue (Table S3). Despite similar rates of GWAS locus tagging by eQTLs, GENCODE-informed annotations identified approximately 2.5 times as many associations as assembly-derived annotations (Fig. 4a). This ratio was highest in breast tissue (5.04) and fibroblasts (2.30) but was approximately equal in tumor. Importantly, this difference in the number of prioritized hits does not reflect weaker association signals with long-read-informed analyses. Among significant associations, the strength of colocalization signals was comparable across annotations (median posterior probability of shared causal variant: 0.98 for long-read, 0.98 for combined, 0.97 for GENCODE). Similarly, the magnitude of significant TWAS effect sizes did not meaningfully differ (median 4.90 for GENCODE, 4.79 for long-read, and 4.87 for combined). Thus, fewer prioritized isoforms from long-read annotations do not reflect weaker statistical support.

**Figure 4.**
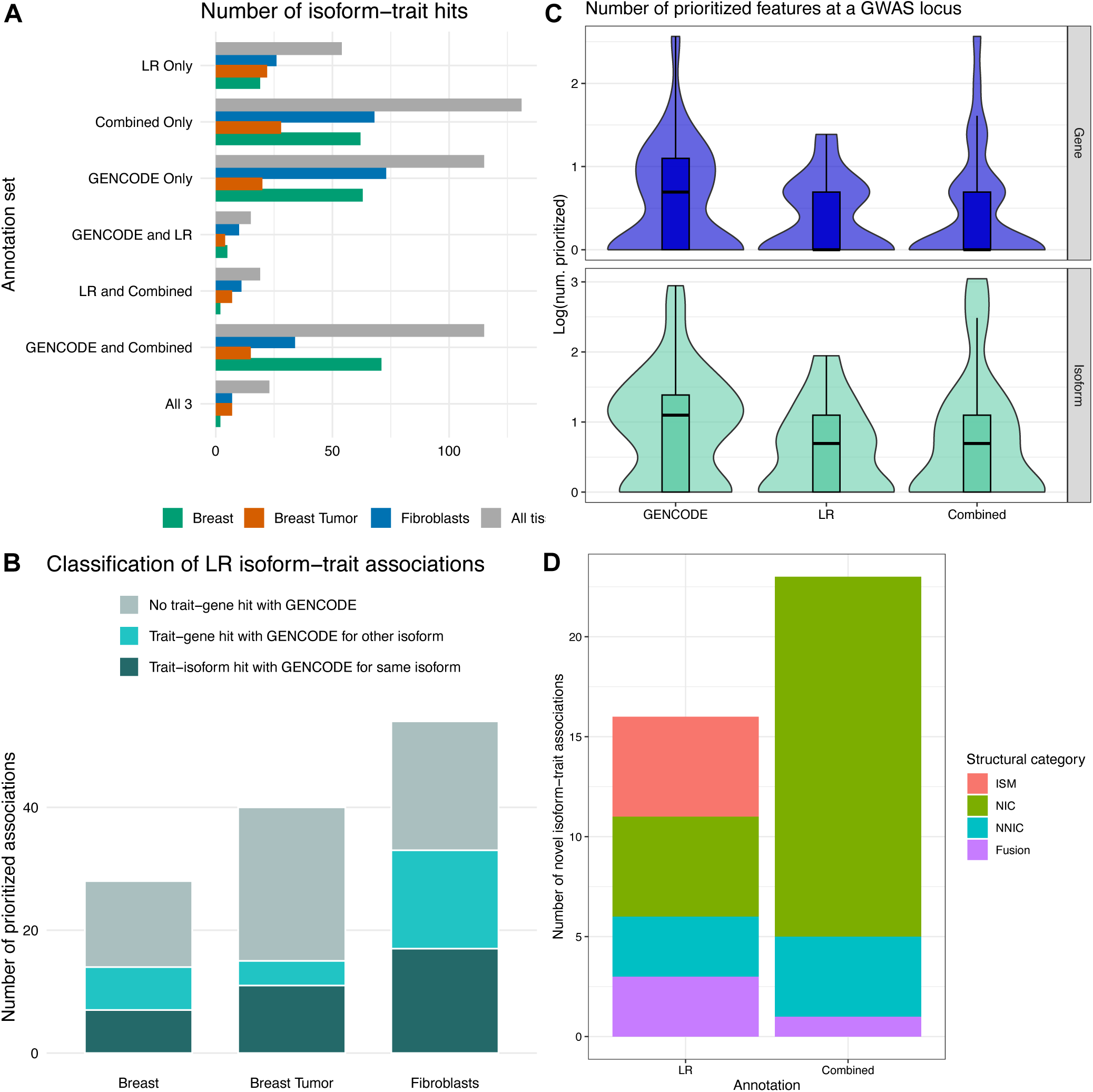
(**A**) Plots show the number of isoform-trait associations prioritized by both TWAS and colocalization in each annotation set for each tissue type. Unique hits aggregated across all tissues are shown in gray. Sets representing intersections of two annotations (e.g., GENCODE and LR) indicate associations identified only in those two annotations and not the third. (**B**) Bar height indicates the number of isoform-trait associations identified using long-read-derived transcriptome annotations by tissue. Bar fill indicates the proportion of associations with a corresponding significant isoform-level association for the same isoform and trait in GENCODE (dark blue), with a corresponding isoform-trait hit in GENCODE but for a different isoform of the gene (turquoise), and with no hits in GENCODE for that trait for any isoforms of the gene (pale blue). (**C**) Boxplots show the distribution of the number of prioritized features (top: genes, bottom: isoforms) within 1Mb of a GWAS risk locus by annotation aggregated across tissues. (**D**) Bar heights mark the number of prioritized isoform-trait associations aggregated across tissues for novel, unannotated isoforms only in the LR (left) and the combined annotations (right). Bar fill shows the distribution of the structural category of these novel isoforms. Abbreviations: TWAS, transcriptome-wide association study; LR, long-read; ISM, incomplete-splice match; NIC, novel in catalog; NNIC, novel not in catalog; GWAS, genome-wide association study.

Just as many of the eIsoforms tagging the same GWAS locus were largely discordant, annotation-dependent variability in isoform prioritization was substantial. Of the 556 colocalization and TWAS-prioritized associations, 381 (69%) were unique to a single annotation within a tissue. Additionally, for each phenotype-tissue pair harboring at least one signal, the number of annotation-unique associations consistently exceeded the number identified by two or more annotations (Fig. S9). Notably, this variability occurred despite the high similarity between GENCODE and combined annotations, indicating that even a modest contribution of novel isoforms can substantially alter isoform-level inference. These patterns were even more pronounced for the less common breast cancer subtypes (luminal B, HER2-enriched, luminal B/HER2-, triple negative). Of these 57 tissue-trait-isoform associations, 47 (82.5%) of prioritized hits were annotation-unique.

Many isoform-trait associations identified only with long-read annotations might be expected to also appear with GENCODE, though assigned to a different but structurally similar isoform from the same gene. However, we found that many long-read-prioritized associations lacked any corresponding signal with GENCODE. Specifically, in fibroblasts, healthy breast tissue, and tumor, 38.8%, 50.0%, and 62.5% of long-read-prioritized isoform-trait associations, respectively, showed neither a significant trait association for the same isoform in GENCODE nor any significant association for another isoform of the corresponding gene (Fig. 4b).

Conversely, when examining associations detected only with GENCODE, only 13.8% (healthy breast), 56.2% (breast tumor) and 43.0% (fibroblasts) of unique isoforms prioritized by GENCODE were present in the corresponding long-read annotations. This suggests that up to half of regulatory hypotheses derived from tissue-agnostic annotations may involve transcripts that are weakly expressed or possibly absent from the relevant tissue.

Next, we restricted analyses to prioritized associations within 1MB of independent breast cancer risk GWAS loci. While long-read annotations yielded significantly fewer associations overall, this also reduced the number of candidate isoforms per GWAS locus (Fig. 4c). Of note, GENCODE prioritized up to 19 candidate isoforms (median 3) and 13 candidate genes (median 2) at a GWAS locus, while long-read annotations prioritized at most 7 isoforms (median 2) and 4 genes (median 1) at those same loci. Furthermore, at these shared loci, we observed considerable variation in the top isoform prioritized by colocalization at each locus across all annotation pairs (Fig. S10). For example, in fibroblasts for our two most well-powered phenotypes (overall breast cancer and luminal A-like cancer), GENCODE and combined annotations prioritized the same isoform approximately half of the time while GENCODE- and long-read-prioritized isoforms agreed only approximately 30% of the time. Variability was even greater in breast tumor analyses, although interpretation was limited by the much smaller number of overall association signals in that tissue.

Long-read assemblies also have the distinct advantage of enabling direct evaluation of novel, previously unannotated transcript structures. While both combined and long-read-only annotations identified significant associations for novel isoforms (23 and 16, respectively), those from the long-read annotations exhibited greater structural diversity (Fig. 4d). These included five hits for transcripts categorized as incomplete-splice matches, while isoforms identified by the combined annotations were largely novel-in-catalog transcripts.

### Long-read annotations prioritize new regulatory features

*MARK1* (1q41) illustrates a high-confidence isoform-trait association detected only with long-read annotations, with no corresponding TWAS or colocalization signal identified under GENCODE. Only the fibroblast long-read annotation identified a putative driver isoform associated with both overall breast cancer risk and luminal A-like cancer. *MARK1* encodes a serine/threonine protein kinase involved in the regulation of microtubule dynamics and cell polarity through the phosphorylation of microtubule-associated proteins^34^. This gene has been recently implicated in the prediction of treatment response in HER2-positive breast cancer tumors to targeted therapy^35^ and has also been studied as a target of dysregulated miRNAs in cervical cancer^36^.

GENCODE version 45 catalogs 18 *MARK1* transcript-isoforms (Fig. 5a). However, when we quantified the short-read fibroblast samples using GENCODE, only two of these isoforms passed the minimum expression threshold (TPM>0.1 in ≥25% of samples) and were retained for analysis (Fig. 5b). By contrast, when quantifying the same samples using the long-read annotation, two isoforms also passed the expression threshold, but these were completely distinct from the two retained under GENCODE. This gene precisely illustrates how two underlying annotated isoform spaces used for expression quantification may partially overlap, but the isoform sets surviving standard post-quantification quality control can be completely distinct.

**Figure 5.**
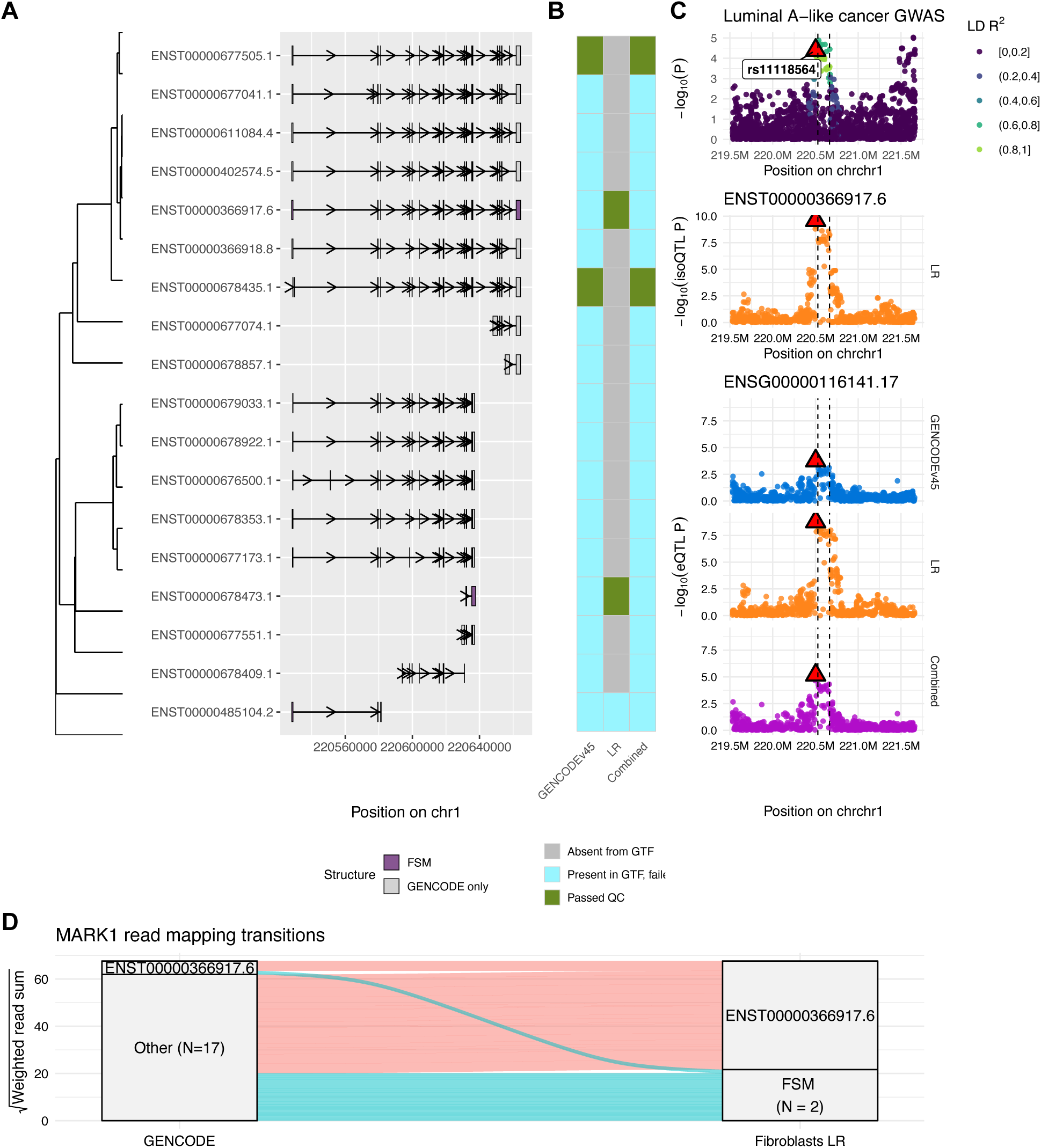
(**A**) Exon-intron structures of *MARK1* transcript isoforms on chromosome 1. Exons are colored by isoform presence in the fibroblast long-read assembly (FSM, purple) or GENCODE only (gray). (**B**) Isoform detection and QC in fibroblasts: gray indicates isoforms not detected in the long-read assembly, blue indicates detected isoforms that failed quality control filtering, and green indicates isoforms that passed. (**C**) Regulatory signals in the *MARK1* locus. Top panel: −log10(GWAS P-value) for variants within 1 Mb of the gene. The lead eQTL variant for the LR-prioritized isoform (rs11118564) is marked by a red triangle. Variant colors reflect LD R² with this lead variant in LR fibroblasts. Middle panel: eQTL signal for the prioritized isoform (ENST00000366917.6) in the long-read annotation. Bottom three panels: gene-level eQTL signals across all three transcript annotations in fibroblasts. (**D**) Short-read alignment transitions to *MARK1* transcripts in five randomly selected GTEx fibroblast samples. Left: alignments to GENCODE annotation; right: alignments to LR fibroblast annotation. All transcripts except the LR-prioritized isoform are collapsed for ease of visualization. Abbreviations: FSM, full-splice match; LR, long-read; LD, linkage disequilibrium; GWAS, genome-wide association study; eQTL, expression quantitative trait loci; isoQTL, isoform eQTL; GTF, gene transfer format file; QC, quality control.

Across GENCODE, long-read, and combined annotations, we observed only one significant colocalization between GWAS and isoQTL signals at this locus (Fig. 5c). This signal corresponded to ENST00000366917 and was driven by lead variant rs11118564 in the long-read annotation. Gene-level eQTLs were significant only under the long-read annotation and attenuated when quantified using GENCODE or the combined references. Mapping short-read alignments from five randomly selected GTEx fibroblast samples using STAR showed that reads supporting this driver isoform were dispersed across multiple structurally similar transcripts when quantified against GENCODE (Fig. 5d), thereby diluting isoform-specific regulatory signal.

While *MARK1* illustrates how long-read annotations can refine regulatory signals within a fully annotated transcript space, long-read assemblies also uniquely enable the discovery and functional evaluation of novel, previously unannotated transcript structures. One example is *NUP107* (12q15, Fig. 6). *NUP107* encodes a core component of the nuclear pore complex, and its associated subcomplex is essential for mRNA export and cell differentiation^37^. While not a canonical cancer susceptibility gene, *NUP107* transcripts were prioritized by both colocalization and TWAS in fibroblasts under GENCODE (ENST00000378905.6) and long-read annotations (lead SNP rs7976687). However, the long-read prioritized isoform is a novel-in-catalog isoform distinguished from reference transcripts by an exon composed of previously annotated splice junctions at chr12:68713730-68715740 (Fig. 6a). Integrated 18-state chromatin learning data from fibroblasts in the ROADMAP Epigenomics Project^38^ showed that the start of this exon coincided with a predicted enhancer (Fig. 6b). This observation supports a model in which exon inclusion may be regulated by chromatin looping with the promoter or through effects on transcriptional elongation and splicing dynamics.

**Figure 6.**
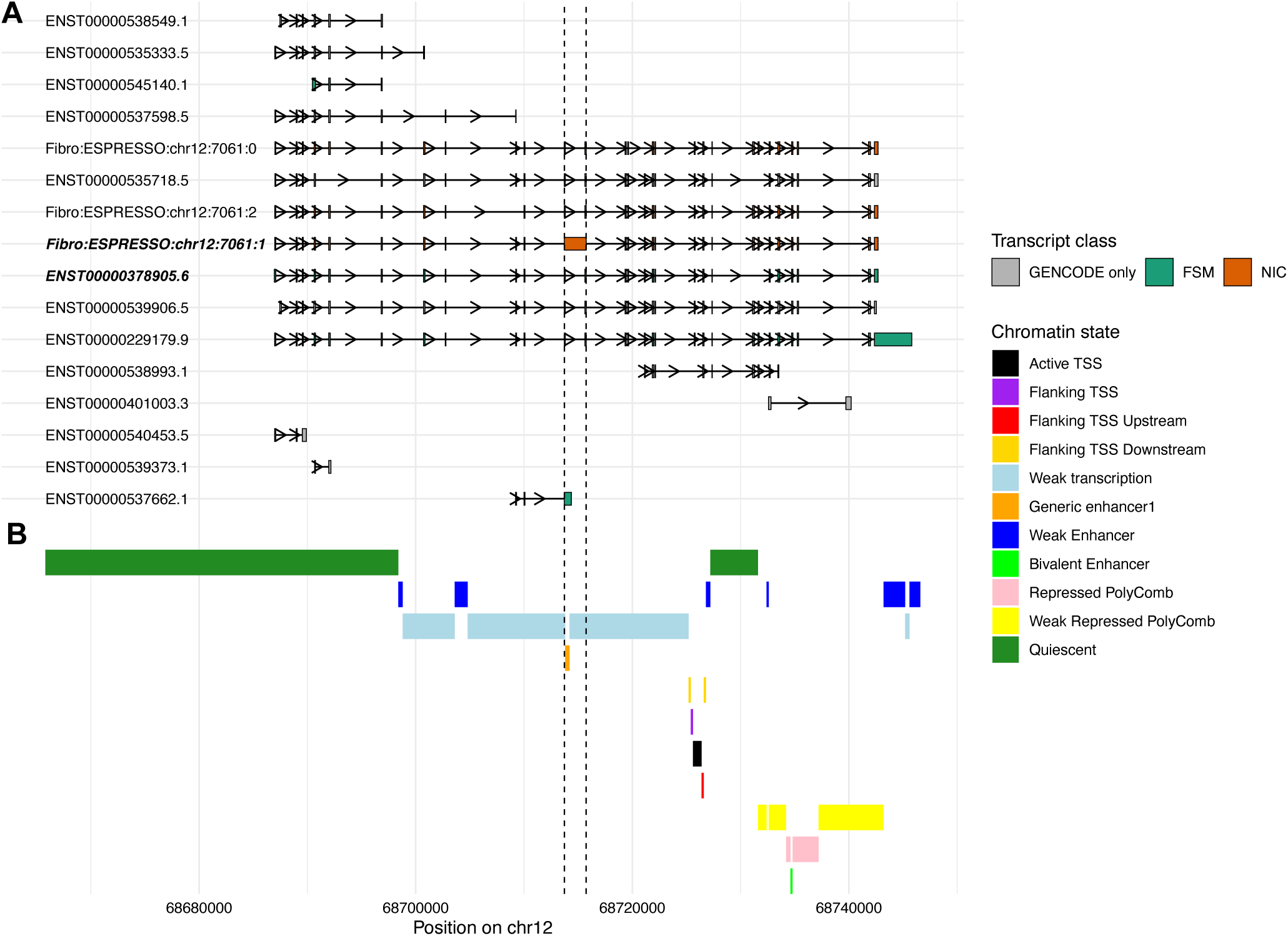
(**A**) Exon-intron structures of *NUP107* transcript isoforms on chromosome 12. Exons are colored by isoform presence and structural classification in the fibroblast long-read assembly: FSM (green), GENCODE only (gray), and NIC (orange). The prioritized novel isoform assembled with ESPRESSO and the GENCODE-prioritized isoform of this gene are bolded. (**B**) Chromatin state landscape across the *NUP107* locus. Genomic regions are annotated using the 18-state ChromHMM model from ROADMAP Epigenomics lung fibroblasts. Colors indicate predicted functional states, including active TSS, weak enhancers, generic enhancers, quiescent regions, and other functional categories. This track highlights the local regulatory context overlapping the LR-prioritized isoform. Abbreviations: FSM, full splice match; NIC, novel in catalog; TSS, transcription start site; LR, long-read.

## DISCUSSION

In this study, we identified 556 unique isoform-level associations with breast cancer risk using three short-read RNA-seq datasets from malignant and normal breast tissues and cultured fibroblasts. We found striking differences in the association patterns depending on the transcript annotations used for expression quantification. These findings suggest that transcriptome annotations should be treated not merely as default inputs in integrative analyses of gene regulation, but rather as biologically meaningful choices that shape downstream inference. Tissue-specific long-read assemblies (1) substantially reduce the isoform space, focusing the hypothesis space for causal isoforms in the relevant tissue context, (2) avoid spurious associations with tissue-irrelevant transcripts, and (3) improve the specificity of inferred regulatory mechanisms at GWAS loci. By prioritizing empirically supported isoforms, long-read-informed annotations can help resolve ambiguity at the isoform level, the most fundamental unit of transcription that remains unexplored in most integrative analyses of complex traits.

TWAS have come under recent scrutiny for high false positive rates, high false sign rates, and limited replicability^33,39–41^. While the polygenicity of complex traits has been proposed as one major driver of these challenges, our current and previous findings suggest that variability in transcriptome annotations and quantification methods may also contribute substantially^33^. Standard transcriptomic references such as GENCODE and Ensembl are largely tissue-agnostic and catalog hundreds of thousands of transcripts spanning protein-coding genes, long noncoding RNAs, pseudogenes, and other biotypes. This expansive isoform space continues to grow with each release and may increase the likelihood of read misassignment to lowly expressed or tissue-irrelevant transcripts. In turn, this can dilute true regulatory signals and ultimately obscure biologically relevant mechanisms. Additional critical strides must be made to better understand how annotation choice shapes downstream genetic inference.

One such critical next step is to accurately quantify how uncertainty arising from overpopulated transcript references impacts statistical power and type I error of eQTL discovery, TWAS, and colocalization studies. Our results directly motivate the development of improved simulation frameworks to address this question. Existing tools for QTL simulation rely on simplified distributional assumptions that do not capture the complexity of isoform-level RNA-seq data. More realistic approaches that generate raw sequencing reads, model multiple modes of genetic regulation, and incorporate modern library preparation protocols will be needed to systematically evaluate these annotation-dependent biases in downstream analyses.

While our results suggest that re-quantification using tissue-matched long-read-informed annotations provides the most precise framework for regulatory inference, this approach may not always be feasible for large legacy datasets. These constraints motivate the development of methods that preserve the benefits of annotation-aware inference while reducing the need for full RNA-seq reprocessing. Moving beyond simple associations toward causal resolution of driver isoforms will likely require isoform-level fine-mapping frameworks that jointly model the effects of individual transcripts, total gene expression, and nearby regulatory variants. We believe incorporating biologically informed priors derived from long-read evidence (e.g., structural support, splice junction validation) may further improve causal inference while reducing the need for large-scale resequencing efforts.

Long-read panels remain limited by sample size, high sequencing costs, and scattered representation across tissue contexts. In this study, healthy breast tissue posed a particular challenge despite its importance for modeling germline susceptibility to breast cancer and its status as the gold-standard context in existing breast cancer TWAS. Limited long-read sample sizes can lead to incomplete transcript capture and reduced sensitivity for low-abundance isoforms, which in turn, can severely limit discovery. As publicly available long-read resources expand, combining samples across independent and diverse short-read cohorts for a given tissue may help improve power while preserving tissue specificity. Despite these challenges, we are optimistic that ongoing technological advances and growing long-read catalogs will likely substantially mitigate these concerns.

## METHODS

### Long-read transcriptome assembly construction

We obtained raw long-read RNA sequencing FASTQ files via the European Genome Archive database (accession number EGAS00001004819) from Veiga et al.^18^, who generated PacBio single-molecule reads from a total of 30 breast tissue samples. These included 4 healthy breast samples (two primary tissues and two from cell lines HMEC and HS578BST) and 26 breast cancer samples comprising 13 primary breast tumor biopsies (covering hormone receptor-positive, HER2+, and triple-negative subtypes), 9 patient-derived xenograft tumors, and 4 breast cancer cell lines (HS578T, MCF-7, MDA-MB-231, MDA-MB-468). Detailed sample characteristics (subtype, origin, receptor status, patient demographics) are available in the supplement of the original publication, where applicable^18^. We also obtained 22 long-read RNA sequencing samples from cell-cultured fibroblasts and one non-malignant breast mammary tissue sample from GTEx V9^42^. Fibroblast samples were derived from skin biopsies from the lower leg, and cell lines were cultured under standard conditions with a subset undergoing *PTBP1* siRNA knockdown, high-quality RNA extraction, poly(A) selection, cDNA library preparation, and Oxford Nanopore long-read sequencing according to manufacturer protocols described elsewhere^43^. These raw long-read data served as input for the transcriptome assembly and quantification procedures described below.

We converted raw FASTQ files to unmapped BAM files with samtools^44^ import and processed as in Veiga et al^18^ with the IsoSeq workflow^45^. Briefly, we used lima to remove New England Biolabs and Clontech adapters and to orient the reads. We used IsoSeq refine to filter out reads without poly(A) tails and then trim poly(A) tails and concatemers. We then aligned the full-length non-concatemer poly(A) reads to the human genome assembly analysis set hg38 (GenBank assembly 000001405.15, no ALT contigs) using minimap2^46^ with a k-mer length of 14, minimizer window size of 4, and in splice-aware mode with –splice-flank=no to relax assumptions about bases adjacent to splice junction donors and acceptors. We filtered secondary alignments and coordinate sorted primary alignments with samtools^44^.

For the healthy breast, tumor, and fibroblast samples separately, we merged the filtered and sorted alignments from chromosomes 1-22, X, Y, and M across all samples into one BAM file. This served as input for *de novo* transcript assembly using ESPRESSO^26^. This method has performed well in maximizing discovery while minimizing false positives in error-prone long-read RNA-seq data^26^. We then used SQANTI^27^ to characterize the *de novo* assemblies and classify isoforms into the following structural categories relative to GENCODE v45 transcripts: full splice matches (FSM) with a perfect structural match to an annotated isoform in GENCODE, incomplete splice matches (ISM) with some splice junctions matching reference transcripts, novel in catalog (NIC) with new combinations of annotated splice junctions in reference transcripts, novel not in catalog (NNC) with novel donors and/or receptors, and other classes such as fusion products and antisense transcripts^47^.

We implemented custom filtering protocols to define high-confidence ESPRESSO transcripts with integrated orthogonal evidence including short-read splicing junction support, proximity to external transcription start and termination site (TSS, TTS) annotations, and sequence-based artifact filtering. Orthogonal sources included DNase-seq peaks from ENCODE^48^, 3’ and sequencing alternative polyadenylation site data from PolyASite^49–51^, and CAGE-seq peaks from PHANTOM5^52^. We provide full accessions in Table S4. We applied filters to novel isoforms to remove likely technical artifacts such as RT switching, intrapriming, and spurious nonsense-mediated decay predictions. We provide descriptions of full quality control criteria elsewhere^53^.

We generated “combined” gene and transcript isoform annotations by integrating high-confidence novel isoforms from our long-read healthy breast, breast tumor, and fibroblast assemblies with the GENCODE reference. We constructed a transcript-to-gene mapping by assigning transcripts to annotated genes or, when absent, to novel gene IDs. We then appended novel genes and transcript isoforms from ESPRESSO absent in GENCODE to the reference. We harmonized merged annotations, resulting in two combined gene transfer format (GTF) files containing both GENCODE genes and high-confidence long-read transcripts for downstream analyses per tissue. We then used MashMap^54^ to construct decoy-aware transcriptome indices for Salmon^55^ for the long-read-derived healthy, tumor, and fibroblast assemblies, the reference GENCODE annotation, and the three combined annotations of GENCODE concatenated with unannotated non-FSM transcripts from the long-read assemblies.

### Genotype processing

#### GTEx V8 dataset

We describe the full protocols for processing GTEx genotypes elsewhere^8^. Briefly, we obtained genotypes from GTEx V8^56^ from whole-genome sequencing data of 838 postmortem donors. We retained only single nucleotide polymorphisms (SNPs) and next filtered out variants with elevated genotype missingness (>5%), variants with minor allele frequency (MAF) <1%, and those showing significant deviation from Hardy-Weinberg equilibrium (P < 10E-6). We additionally removed SNPs with a MAF<1% in the European ancestry subset of the 1000 Genomes Project^57^. We were left with a total of 1,131,422 autosomal variants (GRCh38) for analysis.

#### TCGA dataset

We converted Affymetrix SNP 6.0 CEL files for TCGA breast carcinoma (BRCA) blood-derived normal samples to variant call format (VCF) using a custom R script. We first prepared a probe annotation file mapping SNP probe identifiers to GRCh38 coordinates by obtaining hg19 probe positions from the UCSC snpArrayAffy6 table and lifting them to GRCh38 using the UCSC liftOver chain via the rtracklayer R package, retaining probe identifiers, reference and alternative alleles, and strand information. We then performed genotype calling on each CEL file individually using the Corrected Robust Linear Model with Maximum Likelihood Distance (CRLMM) algorithm, implemented in the crlmm R/Bioconductor package^58^. We recorded CRLMM genotype calls to VCF genotype notation and mapped to the lifted-over GRCh38 probe coordinates. We converted genotype confidence scores returned by CRLMM to Phred-scaled genotype quality (GQ) values using the formula GQ = −10 × log10(1 - *p*), where *p* is the CRLMM confidence score, with values capped at 99. Variants with GQ < 20 were flagged as FAIL in the VCF FILTER field, and we wrote per-sample VCFs with GT and GQ FORMAT fields. Probes that failed to return a genotype call were excluded. We filtered VCFs with bcftools to exclude variants with GQ < 20 or FILTER status of FAIL. We sorted filtered VCFs, compressed them with bgzip, and indexed with tabix. We validated variant calls using GATK ValidateVariants.

We phased filtered VCFs using Eagle2^59^ with the 1000 Genomes Project high-coverage phased reference panel. This panel was prepared by filtering for sites with minor allele count ≥ 2, normalizing multiallelic variants with bcftools norm, removing duplicate sites, and converting to BCF format. We used a GRCh38 genetic map for recombination rate estimation.

We performed genotype imputation using Minimac4 v4.1.6 with the 1kGP high-coverage phased reference panel in MSAV format^60^. We concatenated imputed per-chromosome BCFs into a final genome-wide imputed BCF and indexed. Following imputation, we filtered out variants with imputation R^2^<0.8, those with high missingness (>5%), variants with MAF<1%, and those failing Hardy-Weinberg equilibrium (P < 10E-6). Among individuals with two genotyped samples derived from both blood and solid tissue, we preferentially kept those from blood. We were left with a total of 2,679,319 autosomal variants (GRCh38) for analysis across 1,096 unrelated individuals.

### RNA-seq processing

We first obtained 459 aligned BAM files for breast mammary tissue and 504 files for cultured fibroblast samples from the GTEx project V8 (dbGaP: phs000424.v8.p2) that passed GTEx Analysis Freeze quality control. We used SAMtools^44^ to convert the BAM files to paired-end FASTQ format for quantification via Salmon^55^ (described below) and discarded unpaired and orphaned reads. Next, we used trimmomatic^61^ to quality trim the raw paired-end FASTQ files. For quality-control alignment metrics, we built a STAR genome index from the GRCh38 reference FASTA and GENCODEv45 gene annotation and re-aligned the trimmed reads to generate BAM files. For TCGA, we extracted RNA-seq reads from GDC-aligned BRCA BAM files by converting to paired-end FASTQ format using samtools. We performed adapter trimming and quality filtering with fastp using default parameters. We subsequently retained only primary solid tumor breast cancer samples, excluding metastatic and tumor-adjacent samples.

For both GTEx and TCGA datasets, we used Picard^62^ to collect GC bias, RNA-seq, and alignment summary metrics. We aggregated all results with MultiQC^63^ to produce inputs for downstream hidden covariate estimation.

### Expression quantification

We quantified the GTEx trimmed reads using Salmon with three transcript annotations: GENCODEv45, the long-read *de novo* breast and fibroblast assemblies, and the combined GTFs concatenating unannotated non-FSM transcripts to the reference annotation. We similarly quantified trimmed reads with the GENCODE reference transcriptome, as well as the long-read tumor assembly and the combined annotation of those two. We built Salmon v1.10.2 indices for each reference using a decoy-aware strategy. We used the generateDecoyTranscriptome.sh script (provided with Salmon) to construct a gentrome FASTA by appending the GRCh38 reference genome (GCA_000001405.15, no-alt analysis set) as decoy sequence to the transcript FASTA. We used MashMap for decoy identification and BEDtools for coordinate extraction. We generated Salmon indices from the resulting gentrome with a k-mer size of 31 and decoy sequence identifiers provided via a decoys list. We performed transcript-level quantification with Salmon in mapping-based mode^55^ using automatic library type detection, mapping validation, sequence-specific bias correction, and 50 bootstrap replicates for uncertainty estimation.

We imported isoform-level quantifications into R using tximeta^64^ and removed non-autosomal transcripts. We aggregated isoform-level quantifications to the gene-level using summarizeToGene() from tximeta. We removed lowly expressed transcripts and genes by requiring each to have TPM>0.1 in at least 25% of samples. To address overdispersion arising from read-to-transcript ambiguity, we applied catchSalmon() from the edgeR^65^ package and divided TPM of each isoform by its estimated overdispersion factor. We normalized gene- and transcript-level quantifications using trimmed-method of means (TMM, edgeR^65^) and subsequently transformed them to log scale using the variance-stabilizing transformation implemented in DESeq2^66^. We then removed outlier samples with gene-level WGCNA network connectivity scores below −3^67^.

To visually assess read mapping transitions between GENCODE and the long-read-derived fibroblast annotation, we also performed alignment of reads with STAR in five randomly selected short-read samples.

### eQTL mapping and fine-mapping

Prior to eQTL mapping, we first estimated hidden covariates to control for latent technical variation in expression matrices. We estimated these components using a regularized latent factor model incorporating RNA-seq, GC content, and genome-level quality metrics from Picard as prior covariates^62,68^. We applied this procedure to normalized gene- and isoform-level expression matrices and inferred k latent components for downstream analyses. For each annotation and tissue, we ran this procedure four times assuming k=(15,50,100,200) hidden technical covariates.

We next performed gene- and isoform-level *cis*-eQTL mapping using QTLtools^69^ with the conditional linear regression framework. We first ran permutation-based analysis with 1,000 permutations to inform the nominal p-value thresholds at FDR 5%. This permutation procedure adjusts nominal p-values for the number of variants in the *cis* region and LD within this 1MB window. We corrected expression phenotypes for covariates (age, sex, top 5 genetic principal components to control for population structure, and hidden covariates). We determined the optimal number of hidden covariates for use in eQTL mapping, as well as colocalization and TWAS, as those that maximized eGenes and eIsoform discovery, respectively. Following WGCNA-based outlier removal and restriction to samples with complete genotype, covariate, and sequencing data, we retained 483/483/476 fibroblasts (GENCODE/combined/LR), 360/360/362 healthy breast samples, and 1,083/1,085/1,085 tumors. Differences in sample counts reflect minor annotation-specific variation in WGCNA-based outlier detection. After covariate adjustment, we rank normal transformed each adjusted expression vector. Following this, we performed eQTL fine-mapping using the SuSiE Bayesian variable selection framework^70^. We again used a 1MB window around the gene or isoform to extract genotype data and centered and scaled genotypes. With residualized expression matrices, we fit SuSiE with L=10 single-effect components to estimate posterior inclusion probabilities (PIP) and identify credible sets with 95% minimum coverage.

### BCAC GWAS summary statistics

We obtained summary-level breast cancer GWAS data from the Breast Cancer Association Consortium (BCAC, see Web Resources). The summary statistics for variant-level associations with breast cancer risk and risk of specific breast cancer subtypes were the result of a large multi-study GWAS of women of European genetic ancestry^5^. The overall breast cancer analysis used genotype data from cases (invasive, *in situ*, unknown invasiveness) and controls across 82 BCAC studies that were genotyped using either the iCOGS or OncoArray Illumina genome-wide custom arrays. For this overall analysis, data from 11 other breast cancer GWASs were incorporated. This yielded a total sample size of 133,384 cases and 113,789 controls. The authors estimated SNP-disease associations using standard logistic regression, adjusting for country of origin and top principal components. Results were obtained for iCOGS subjects, OncoArray subjects, and additional GWASs separately and then combined via fixed-effects meta-analysis.

In addition to the summary statistics for the outcome of overall breast cancer risk, this study also published summary statistics for the association of variants with risk of specific intrinsic-like subtypes of breast cancer: luminal A-like cancer, luminal B-like cancer, luminal B/HER2-negative-like cancer, HER2-enriched-like cancer, and triple-negative cancer. These subtype analyses were performed by fitting two-stage polytomous logistic regression models. Full details on these models are described elsewhere^71,72^. Only invasive cases were considered for this analysis, and samples from the 11 additional GWASs were not included due to missing tumor marker information. The final sample for the GWAS subtype analyses included 106,278 cases and 91,477 controls.

Lastly, a meta-analysis was performed combining results from the analysis of triple-negative breast cancer cases from the BCAC and the separate analysis of cases and controls with a pathogenic *BRCA1* variant from the Consortium of Investigators of Modifiers of BRCA1/2 (CIMBA)^73^. The CIMBA participants were also of European ancestry. Authors performed a fixed-effects meta-analysis, combining odds ratio estimates from the BCAC study and hazard ratio estimates from the CIMBA study. As approximately 69% of breast cancer cases in individuals with *BRCA1* mutations are triple negative^74^, our “triple-negative” TWAS results utilized summary statistics from this meta-analysis for greatest power. Quality control, imputation, and ancestry classification protocols for all sets of genotype data used in the study are described separately^1,6,75,76^, and additional details on the statistical methods used in the breast cancer GWAS are provided by Zhang et al.^5^ The full list of all contributing studies to each analysis (overall breast cancer, subtype specific, and CIMBA) can be found in Tables S1-S3 of Zhang et al.^5^ Once we obtained all summary data from these GWASs, we performed liftover to map GWAS variants to GRCh38. Subsequently, we removed any variants with imputation R^2^ below 0.3 when available, strand-ambiguous alleles, and non-biallelic SNPs.

To define independent risk loci across all 6 sets of breast cancer GWAS summary statistics, we first pruned GWAS SNPs to those with P < 5E-8. We then performed LD-based clumping in PLINK^77^ using the 1000 Genomes European reference panel^57^ (R² < 0.05 within 1 MB and secondary threshold P < 1E-4) and retained the lead SNP per locus. This procedure yielded 332 independent breast cancer risk loci.

### Colocalization and isoform-level TWAS

We performed Bayesian colocalization at the gene- and isoform-resolution to test for a single shared genetic variant driver of both expression signals from marginal variant-level linear regression and GWAS summary statistics using the coloc.abf() functionality within coloc with default priors^78^. Next, we utilized the isoTWAS framework to train predictive models of isoform expression using all cis-SNPs within 1MB of each gene body across the six breast cancer phenotypes. Details on these statistical models are provided elsewhere^8^. We selected the optimal model for each isoform with 10-fold cross-validation, and we retained only those features with cross-validated R^2^ > 0.01. These models were then applied in a weighted burden framework to evaluate associations between the genetically-predicted component of isoform expression and cancer risk. To address the multiple testing burden for isoform-level analyses, we implemented a two-stage testing procedure as described previously^8^. For prioritized isoforms, we further evaluated whether SNP-expression weights contributed signal beyond SNP-disease associations observed in the GWAS using a conservative permutation test. We permuted SNP-isoform weights 10,000 times to generate a null distribution of the test statistic and derived empirical p-values. We restricted permutation testing to features with FDR-adjusted P < 0.05 and isoTWAS hits meeting both false discover rate-adjusted P < 0.05 and family-wide error rate-adjusted P < 0.05.

## Data availability

Long-read RNA-seq-derived transcriptome annotations, scripts, and all code used to generate the data and figures presented in this paper are available upon publication at https://github.com/staylorhead/Long_Read_Informed_Breast_Cancer. The GWAS summary statistic data for risk of overall and subtype-specific breast cancers are publicly available and hosted at: https://www.ccge.medschl.cam.ac.uk/breast-cancer-association-consortium-bcac/data-data-access/summary-results. Individual-level datasets from TCGA were obtained via dbGaP accession number phs000178.v11^79^. Individual-level GTEx data were accessed through dbGaP under accession phs000424.v8.p2^56^. LD reference data from the 1000 Genomes Project^57^ were obtained from the International Genome Sample Resource data portal at https://www.internationalgenome.org/data-portal/sample. The GENCODE human reference transcriptome data (version 45) were downloaded from https://www.gencodegenes.org/human/release_45.html.

## Acknowledgements

This work was supported by the National Institutes of Health (R21CA293419, R01CA237541, R01CA264987).

