## Supplemental figures 1-10 for "Improving isoform-level eQTL and integrative genetic analyses of breast cancer risk with long-read RNA transcript assemblies"

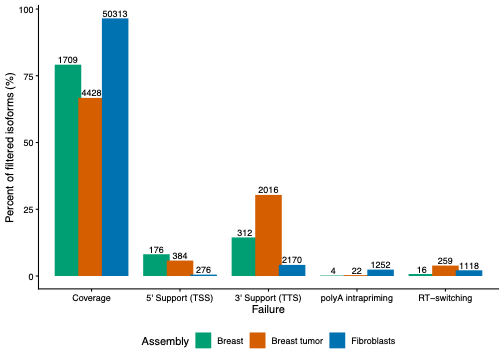

**Figure S1. Filtering reasons for low-confidence isoform calls across long-read assemblies in breast, breast tumor, and fibroblast samples**. Bars indicate the proportion of isoforms excluded by specific quality control criteria among all excluded for a tissue. Coverage filter indicates insufficient short-read or full-length read splice junction support . 5’ support requires the transcription start site (TSS) to be within 10 kb of a CAGE peak or within 1 kb of a reference TSS. 3’ support requires the transcription termination site (TTS) to be within 1 kb of a 3’-seq peak, within 40 bp of a polyA motif, or within 1 kb of a reference TTS. PolyA intrapriming filter, performed only for non-full splice match isoforms, excludes isoforms with <80% adenosine content downstream of the TTS. RT-switching ( for novel not-in-catalog and “other” isoforms) removes isoforms with predicted reverse transcription artifacts at splice junctions.

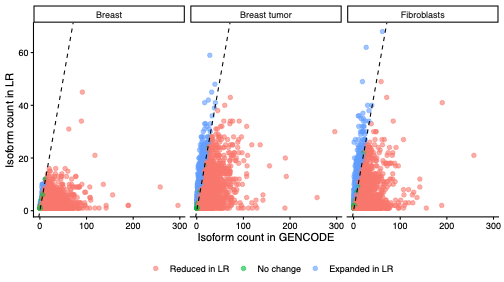

**Figure S2. Comparison of isoform counts per gene in long-read assemblies versus GENCODE**. Scatterplots show the number of isoforms per gene for genes shared between GENCODEv45 and each post-QC long-read assembly (healthy breast, breast tumor, fibroblast). Points are colored by relative change in isoform number: red indicates a decrease in long-read isoforms compared to GENCODE, green indicates no change, and blue indicates an increase.

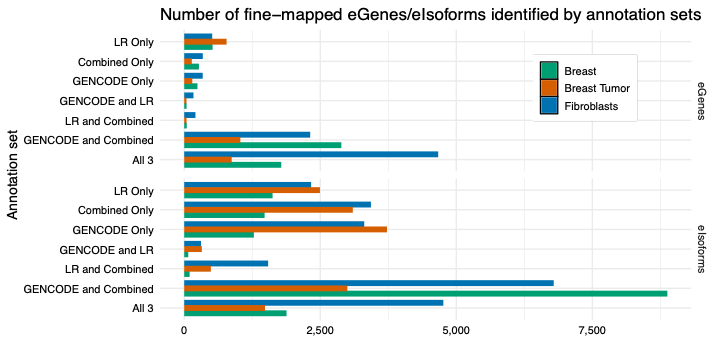

**Figure S3. Overlap of fine-mapped eGenes/eIsoforms across transcriptome annotation sets by tissue**. Plots show the number of eGenes and eIsoforms detected in each annotation set for each tissue type. Sets representing intersections of two annotations (e.g., GENCODE and LR) indicate eGenes identified only in those two annotations and not the third. eGenes were defined using a Bayesian fine-mapping approach with the SuSiE framework. SuSiE was fit with L=10 single-effect components.

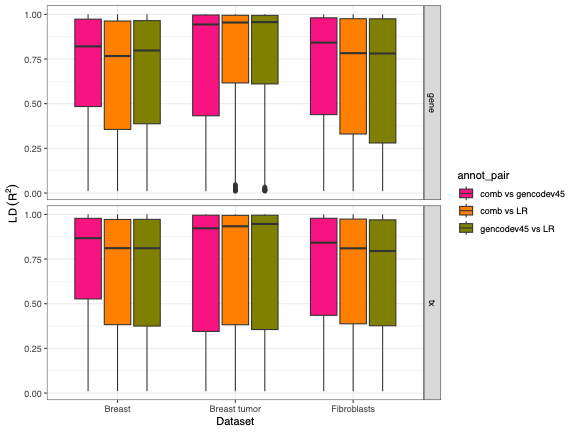

**Figure S4. LD between discordant lead eQTLs for shared eGenes and eIsoforms across annotation pairs**. Boxplots show the distribution of pairwise linkage disequilibrium (LD) r² between lead eQTL variants identified by conditional linear modeling for the same gene or isoform between two annotation sets where the top variant was not the same. LD was calculated using the European reference panel from the 1000 Genomes Project. Results are shown separately for gene- and isoform-level signals and tissue set.

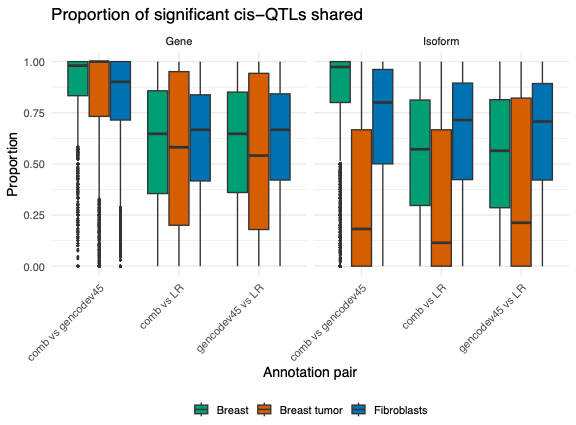

**Figure S5. Proportion of shared significant cis-eQTLs for shared eGenes and eIsoforms across annotation pairs**. Boxplots show, out of all the eQTLs identified for a common eGene/eIsoform across two annotations for a given tissue, what proportion are shared. eQTL variants were identified by conditional linear modeling. Results are shown separately for gene- and isoform-level signals and tissue set.

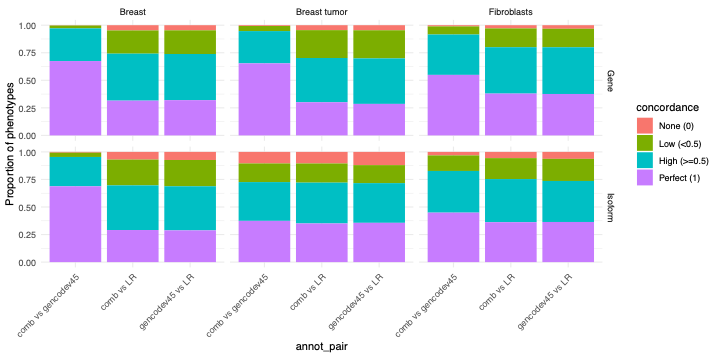

**Figure S6. Overlap of fine-mapped credible set variants across annotation pairs for shared eGenes and eIsoforms**. Stacked barplots show the proportion of shared eFeatures whose 95% credible sets (CS) exhibit no, low, high, or perfect concordance between annotation pairs. Concordance was calculated as the number of shared CS variants divided by the total number of unique CS variants identified across both annotations.

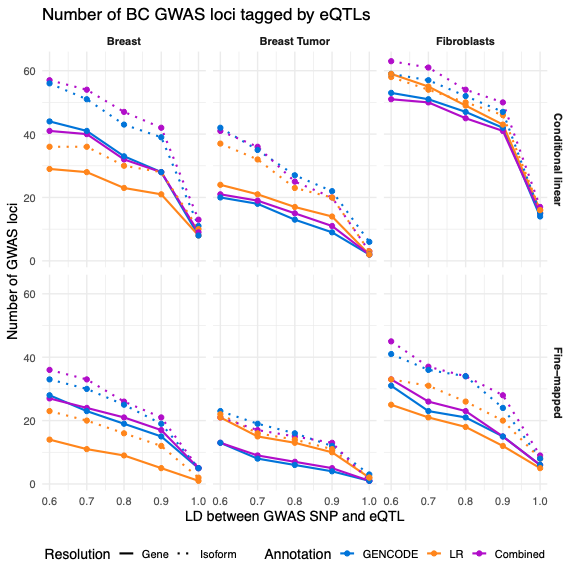

**Figure S7. Gene- and isoform-level eQTL tagging of independent breast cancer GWAS loci by LD threshold.** A tagged locus indicates an eQTL is within 1Mb and LD R^2^ exceeding some value with the GWAS risk variant. Solid lines indicate gene-level eQTLs, and dashed lines indicate transcript-level eQTLs. eQTLs were defined either using conditional linear regression (top row) or Bayesian fine-mapping with SuSiE (bottom row).

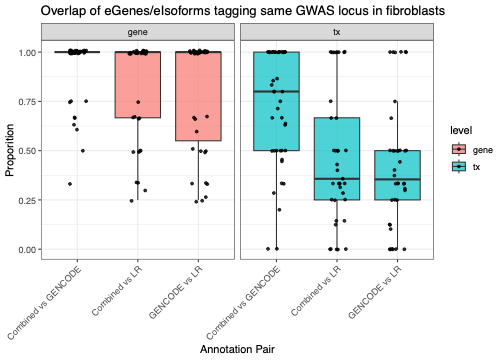

**Figure S8. Proportion of shared regulatory features underlying concordant GWAS locus tagging across annotations in fibroblasts**. Boxplots show, for each annotation pair, the proportion of parent regulatory features shared when gene-level or isoform-level eQTLs tagged the same GWAS locus in fibroblasts. For each shared locus, concordance was defined as the number of shared eGenes or eIsoforms divided by the total number of unique regulatory features identified in either annotation that tagged that locus. eQTLs were identified using conditional linear regression.

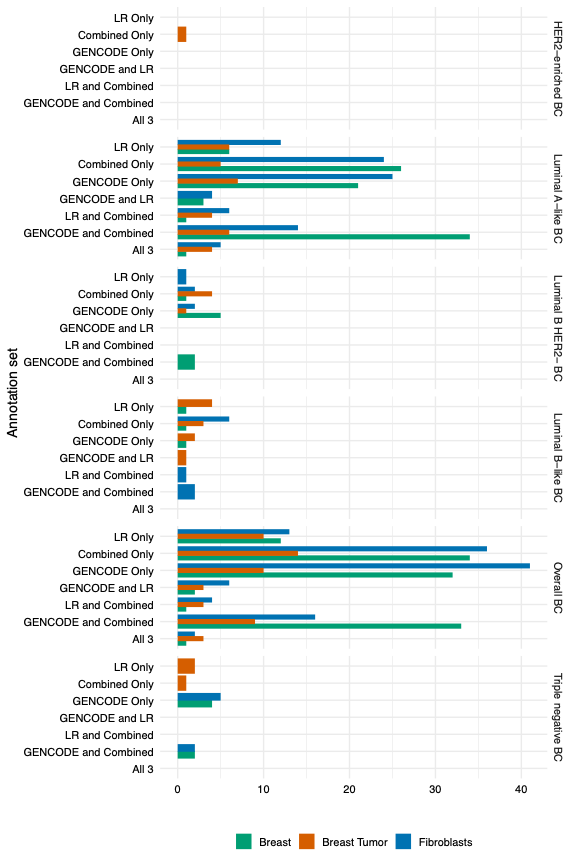

**Figure S9**. **Significant Isoform-trait associations from TWAS and colocalization by breast cancer phenotype.** Plots show the number of isoform-trait associations prioritized by both TWAS and colocalization in each annotation set for each tissue type (fill color). Horizontal panels reflec the six breast cancer phenotypes tested. Sets representing intersections of two annotations (e.g., GENCODE and LR) indicate associations identified only in those two annotations and not the third.

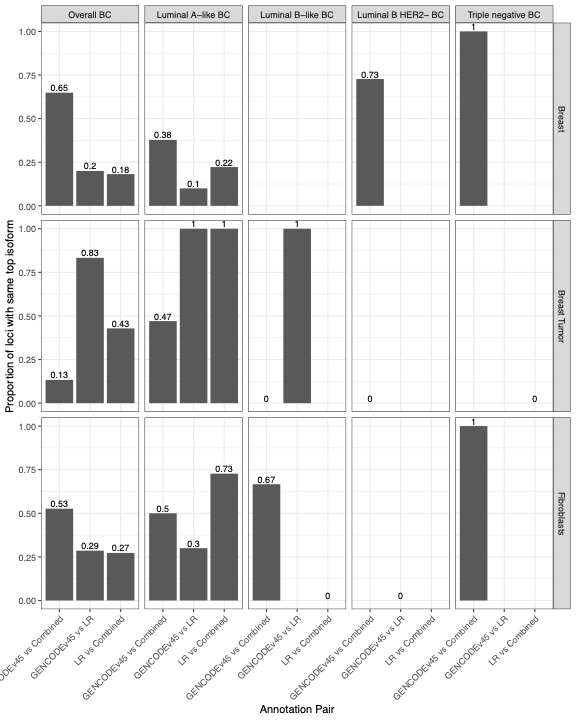

**Figure S10. Concordance of top prioritized isoforms at GWAS loci across annotations.**

For each independent breast cancer GWAS risk locus, we identified significant isoform–trait associations within 1 Mb that were supported by both TWAS and colocalization. Among loci with significant nearby associations detected in more than one transcript annotation for a given tissue and breast cancer phenotype, we calculated the proportion of loci at which the top-prioritized isoform (ranked by colocalization posterior probability, PP.H4) was the same across annotations.
